# Insight into dynamics of APOBEC3G protein in complexes with DNA assessed by high speed AFM

**DOI:** 10.1101/581793

**Authors:** Yangang Pan, Luda S. Shlyakhtenko, Yuri L. Lyubchenko

## Abstract

APOBEC3G (A3G) is a single-stranded DNA (ssDNA) binding protein that restricts the HIV virus by deamination of dC to dU during reverse transcription of the viral genome. A3G has two zing-binding domains: the N-terminal domain (NTD), which efficiently binds ssDNA, and the C-terminal catalytic domain (CTD), which supports deaminase activity of A3G. Until now, structural information on A3G has lacked, preventing elucidation of the molecular mechanisms underlying its interaction with ssDNA and deaminase activity. We have recently built computational model for the full-length A3G monomer and validated its structure by data obtained from time-lapse High-Speed Atomic Force Microscopy (HS AFM). Here time-lapse HS AFM was applied to directly visualize the structure and dynamics of A3G in complexes with ssDNA. Our results demonstrate a highly dynamic structure of A3G, where two domains of the protein fluctuate between compact globular and extended dumbbell structures. Quantitative analysis of our data revealed a substantial increase in the number of A3G dumbbell structures in the presence of the DNA substrate, suggesting the interaction of A3G with the ssDNA substrate stabilizes this dumbbell structure. Based on these data, we developed a model explaining the interaction of globular and dumbbell structures of A3G with ssDNA and suggested a possible role of the dumbbell structure in A3G function.

---

APOBEC3G protein (A3G) belongs to a family of cytidine deaminases^1-3^ with the innate ability to block many retroviruses, including HIV-1 infection in the absence of virion infectivity factor (VIF).^4,5^ A3G was the first and most functionally characterized enzyme.^6^ It was shown that A3G efficiently binds ssDNA and restricts retroviruses with deamination-dependent and deamination-independent restriction pathways.^1,7-12^ A3G has two domains with Z-dependent motifs: the C terminal domain (CTD), which is the catalytically active, and the N-terminal domain (NTD), which is responsible for ssDNA binding.^13^ Both domains contribute to the anti-retroviral activity during viral replication cycle.^14,15^ Attempts to reveal the structure of A3G using traditional methods such as x-ray crystallography and NMR have proved unsuccessful due to the inherent property of A3G to self-assemble into oligomers of various sizes, even at nanomolar concentrations.^2,16-20^ Up to date there is lack of high-resolution atomic structure of full-length A3G; however, structures for individual domains^21-25^ as well as CTD and NTD in complex with ssDNA are available.^26,27^ Based on X-ray crystallography and NMR spectroscopy data for individual domains, we recently^28^ built a computer model for full-length monomeric A3G. The model revealed the dynamics of A3G when two domains change their relative orientation and the protein transforms from a compact globular structure into an extended dumbbell structure. This model was validated by time-lapse high-speed AFM (HS-AFM) and enabled the direct observance of the transition between the globular and dumbbell structures of A3G. Importantly, the ratio between the two structures of A3G obtained from these experiments coincided with simulations, which provide additional validation for the simulated model of the monomeric, full-length structure and dynamics of A3G.

Here the HS-AFM methodology^29-32^ was utilized to visualize the dynamics of monomeric A3G in complex with ssDNA. To unambiguously identify the A3G-DNA complexes, a hybrid-DNA approach^17,28,33,34^ was employed, and different types of DNA substrates were used to reveal the intramolecular dynamics of A3G. It was demonstrated that A3G forms complexes with ssDNA either in compact globular and/or dumbbell structures, but the population of the dumbbell structures of A3G considerably increased compared to the free protein. A clear dependence was also found for the yield of the dumbbell structures on the length of ssDNA substrate. Interestingly, the number of dumbbell structures increases coincidently with the length of ssDNA substrate. The use of different ssDNA substrates allowed us to observe one of the domains being transiently dissociated from ssDNA, demonstrating a very dynamic behavior of A3G in the presence of the ssDNA substrate. Based on these results, we suggested a model to explain the role of the dynamics of A3G in the interaction with ssDNA and form a hypothesis for its role for protein function.

## Results

### Use of DNA substrates in high-speed AFM studies

To examine the structure and dynamics of A3G in complexes with ssDNA, a hybrid-DNA approach was utilized, where ssDNA segments were fused with the DNA duplex, and HS-AFM was applied for unambiguous identification of the A3G ssDNA complexes.^17,28,33,34^ A3G complexes with three different hybrid DNA substrates, as used in this study, are illustrated in Figure S1 A-C. A3G complexes with 69 nt tail ssDNA (A) and 25 nt tail ssDNA (B) show A3G bound to the ssDNA portion next to the dsDNA tag. The A3G complex with 69 nt gap ssDNA (C) illustrates the protein positioned in the ssDNA portion located between DNA duplexes. After assembly of A3G complexes, as described in the Material and Methods section, an aliquot was deposited on APS mica surface for 2 minutes to allow complexes to bind to the surface, followed by rinsing of non-bound complexes and imaging without drying. After the A3G ssDNA complex of interest was selected on the AFM image, continuous frame-by-frame imaging of this complex was performed until A3G dissociated from the ssDNA substrate. The collected frames were then assembled into movies. The corresponding subsections below present the results of data analysis for three different ssDNA substrates in complex with A3G.

### A3G in complex with the 69 nt tail ssDNA substrate

Figure 1 demonstrates the dynamics of A3G in complex with the 69 tail ssDNA substrate, where a few frames were selected from Movie 1. The selected frames demonstrate a highly dynamic behavior of A3G in complex with the 69 nt tail ssDNA substrates, showing both globular and dumbbell structures of A3G. Frame 18 shows the globular conformation of A3G complexed with the 69 nt tail ssDNA. Frame 44 illustrates the transition of A3G from the globular to the dumbbell structure, in which both domains of A3G clearly separate from each other. Frames 56, 57, and 99 demonstrate the fluctuations in the distance between the two domains in the dumbbell structure of A3G, the largest distance shown in frame 99. Later, the domains returned to the globular structure, which is shown in frame 102.

**Figure 1.**
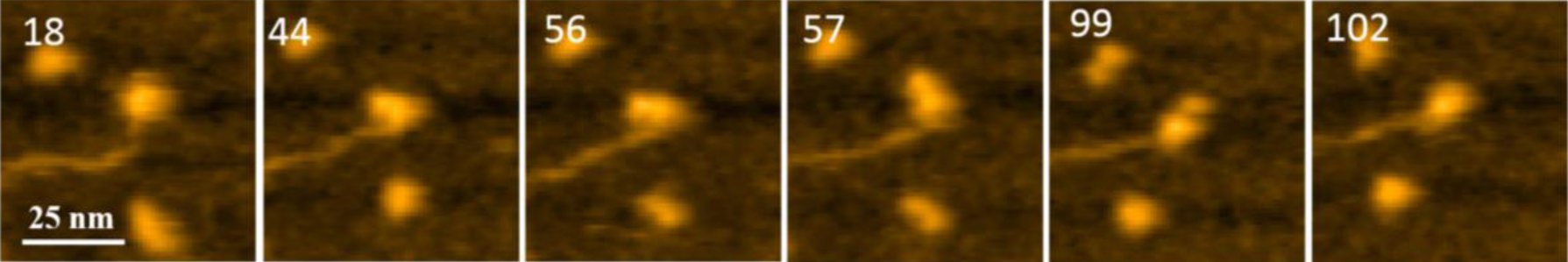
AFM selected frames from Movie 1 illustrating the dynamics of A3G in complex with the 69 nt tail DNA. Frames 18 and 102 show the globular structure of A3G in complex with ssDNA. Frames 44, 56, 57, and 99 represent the dumbbell structure of A3G in complex with ssDNA. The average yield of dumbbell structures is 65%. The scale bar is 25 nm. The scan rate is 398 ms/frame.

The first striking observation for A3G in the complex with the 69 nt tail ssDNA substrate was the high yield of the dumbbell structures. The average yield for the dumbbell structure was 65%, analyzed from 10 separate movies with total ∼600 frames. Note, this yield is four times greater than the yield of A3G dumbbells in free A3G.

For quantitative characterization of the dumbbell and globular structures of A3G in the complex with the 69 nt tail ssDNA, several parameters were used, as shown in Figure 2. For the dumbbell structure of A3G, the cross-sectional feature was selected, as shown in Figure 2A (marked with red line on AFM image). Figure 2B illustrates three parameters, calculated from the cross-section of the dumbbell structure of A3G. The height of each maximum is marked as (h1) for Domain 1 and (h2) for Domain 2; the center-to-center distance is marked as (d) between domains. For the globular structure of A3G, as shown in the AFM image in Figure 2C, the ratio between two orthogonal diameters d1:d2 were used, marked as blue and red lines, respectively. The plot in Figure 2D illustrates measurements for two cross-sections of the globular structure. Figures 3 shows results from data analysis for the dumbbell and globular structures of A3G. Figure 3A shows the dependence of the distance (d) between the two A3G domains on the frame number, calculated for the dumbbell structure of the A3G-69 nt tail ssDNA complexes. These data show a wide range of fluctuation in the distances between two domains, between 3 nm and 8 nm. Figure 3B provides a histogram for the distribution of the distance (d) between the two domains in the dumbbell structure of A3G, and Gaussian fit gives the average distance of d = 5.1 ± 1.0 nm. Figure 3C shows the result for globular A3G in the complex as a dependence of the d1:d2 ratio on the frame number and as the d1:d2 histogram (Figure 3 D). The Gaussian fit to the histogram produces a mean value for the d1:d2 ratio of 1.3 ± 0.2, which resembles data for free A3G.

**Figure 2.**
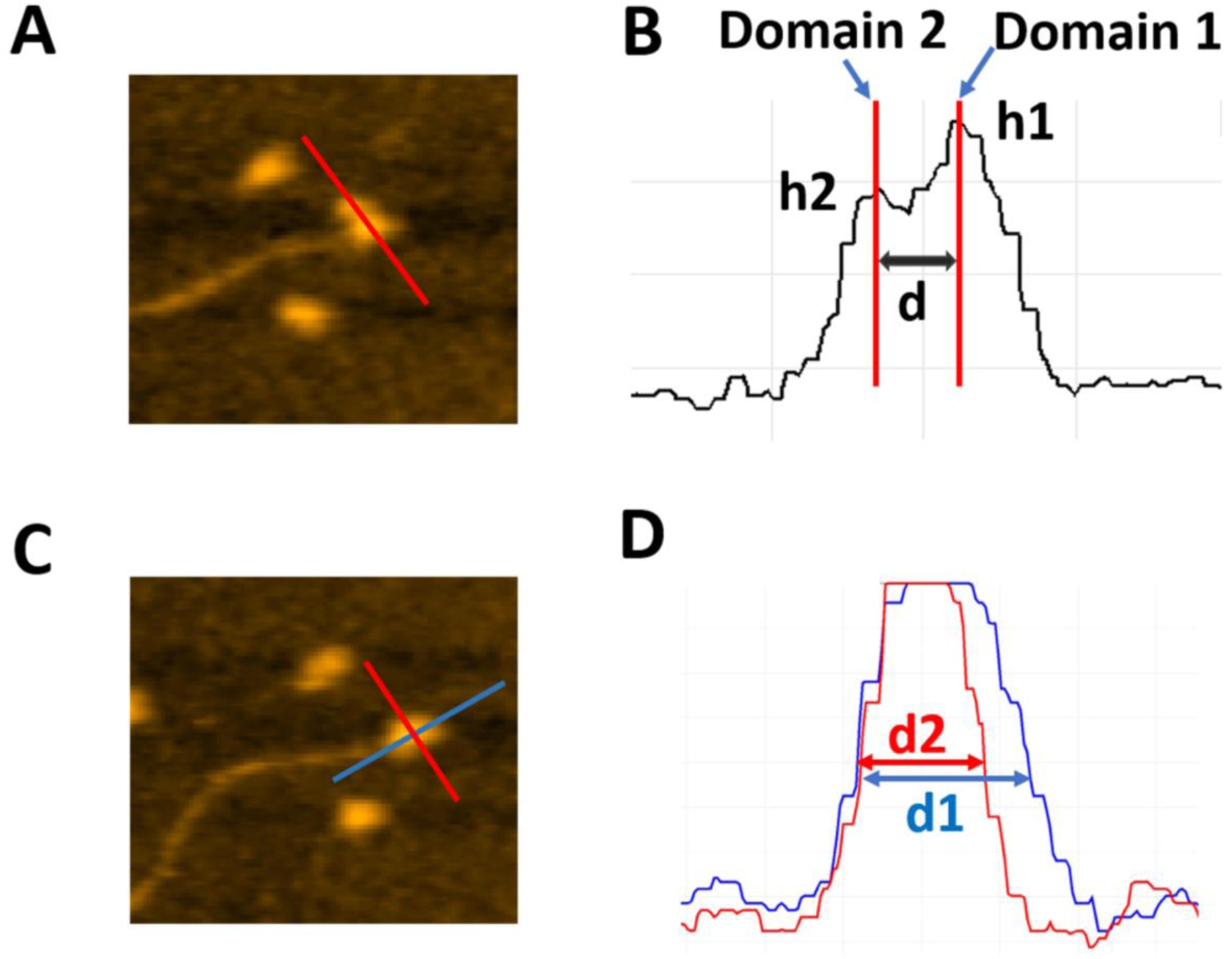
Schematics explaining the analysis of various types of complexes of A3G with DNA. A. AFM image of A3G-69 nt ssDNA complex. The red line shows the cross-section of dumbbell structure for the A3G-ssDNA complex. B. Cross-sectional measurements of the heights of domains (h1) and (h2) and the distance (d) between them. C. AFM image of the dumbbell structure of the A3G-69 nt ssDNA complex. Red and blue lines show the orthogonal cross-sections of the A3G-ssDNA complex. D. Cross-sectional measurements of two orthogonal diameters, d1 and d2.

**Figure 3.**
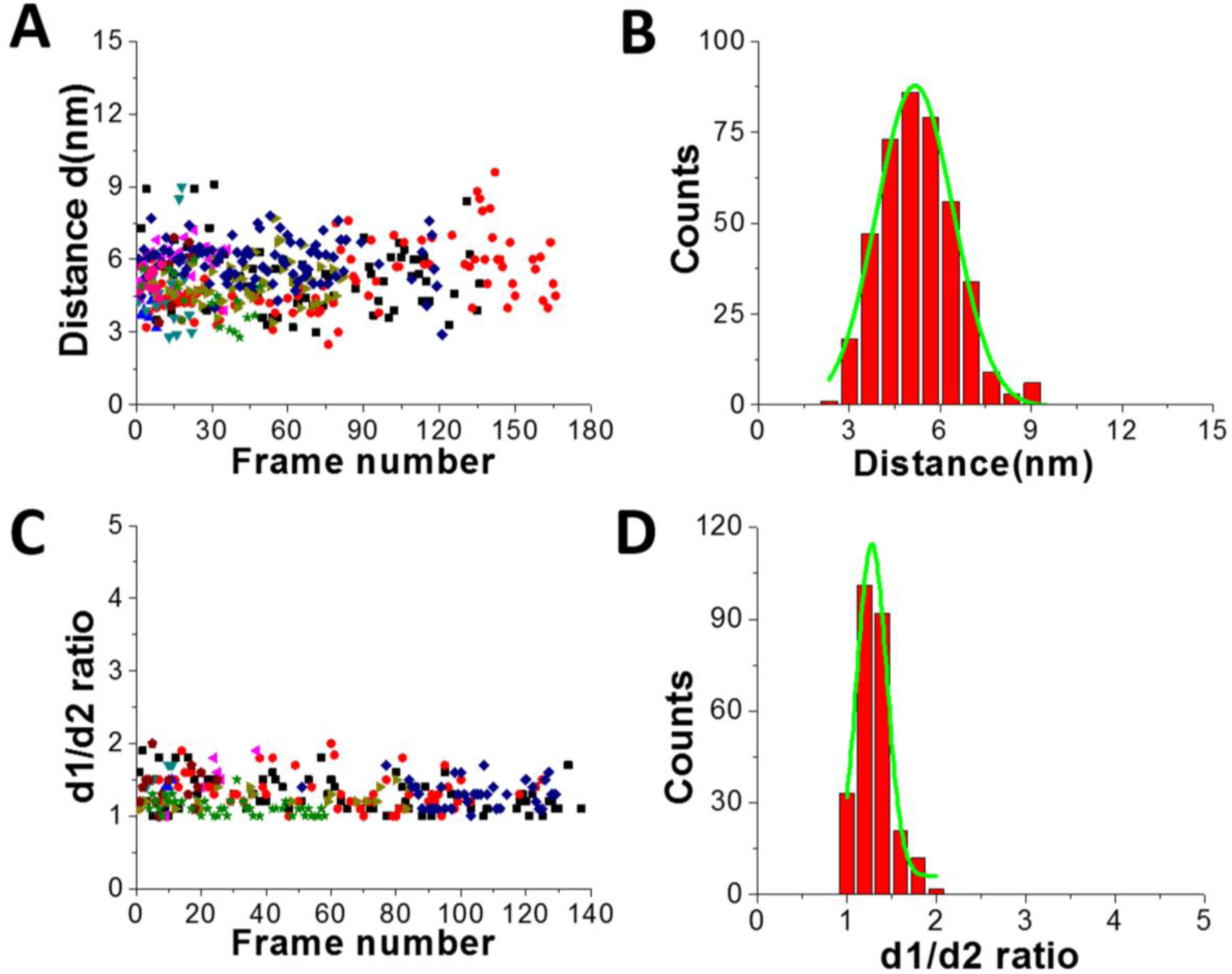
Data analysis for the A3G-69 nt tail ssDNA complex. A. The dependence of distance (d) between Domain 1 and Domain 2 on the frame number for the A3G-69nt tail ssDNA complex. B. The histogram of distances between two domains. The mean value for distances (d) between Domain 1 and Domain 2 together with standard deviation is 5.1 ± 1.0 nm. C. The dependence of the d1:d2 ratio for the globular structure of A3G on frame number for the A3G-69nt tail ssDNA complex. D. The histogram for the d1:d2 ratio. The mean value for d1:d2, with the standard deviation, is 1.3 ± 0.2. The data are the result of analysis of ∼600 frames from 10 separate movies.

Another important parameter, which can be obtained from the HS-AFM data, is the lifetime for the specific structure of A3G in the complex. Figure 4A shows a plot for the dependence of the distance (d) between two domains in the dumbbell structure (right axes, blue) and d1:d2 for the globular structure (left, black) for the A3G-69 nt tail ssDNA complex on the frame number, obtained from one of the movies. Blue dots show changes in the distance (d) between A3G domains in the dumbbell structure, and black triangles represent fluctuations in d1:d2 ratio for the globular structure. Following frame-by-frame transitions between globular and dumbbell structures, the lifetime was calculated for each structure of A3G in the complex. The zoomed portion of the plot in Figure 4A (marked as a red quadrant) is shown in Figure 4B, where several consecutive, uninterrupted frames for the dumbbells characterize their lifetime (blue dots), and likewise, several uninterrupted frames for the globular structure (black triangles) characterize the lifetimes of the globular structure. Figure S2 offers another example of the dynamic behavior of the dumbbell structure of A3G in the DNA complex. The plot in Figure S2 illustrates an example of the long-lived dumbbell structure of A3G in complex with the 69 nt tail ssDNA substrate, with large fluctuations for the distance (d) between the two domains. Analysis of the lifetimes obtained from all assembled movies for the A3G-69 nt tail ssDNA complexes is shown as histograms in Figure 4 for the dumbbells (C) and globular (D) A3G structures. The fit of these histograms with first-order exponential decay gives the lifetime of 0.64 ± 0.03 seconds for dumbbells and 0.39 ± 0.06 seconds for globular structures.

**Figure 4.**
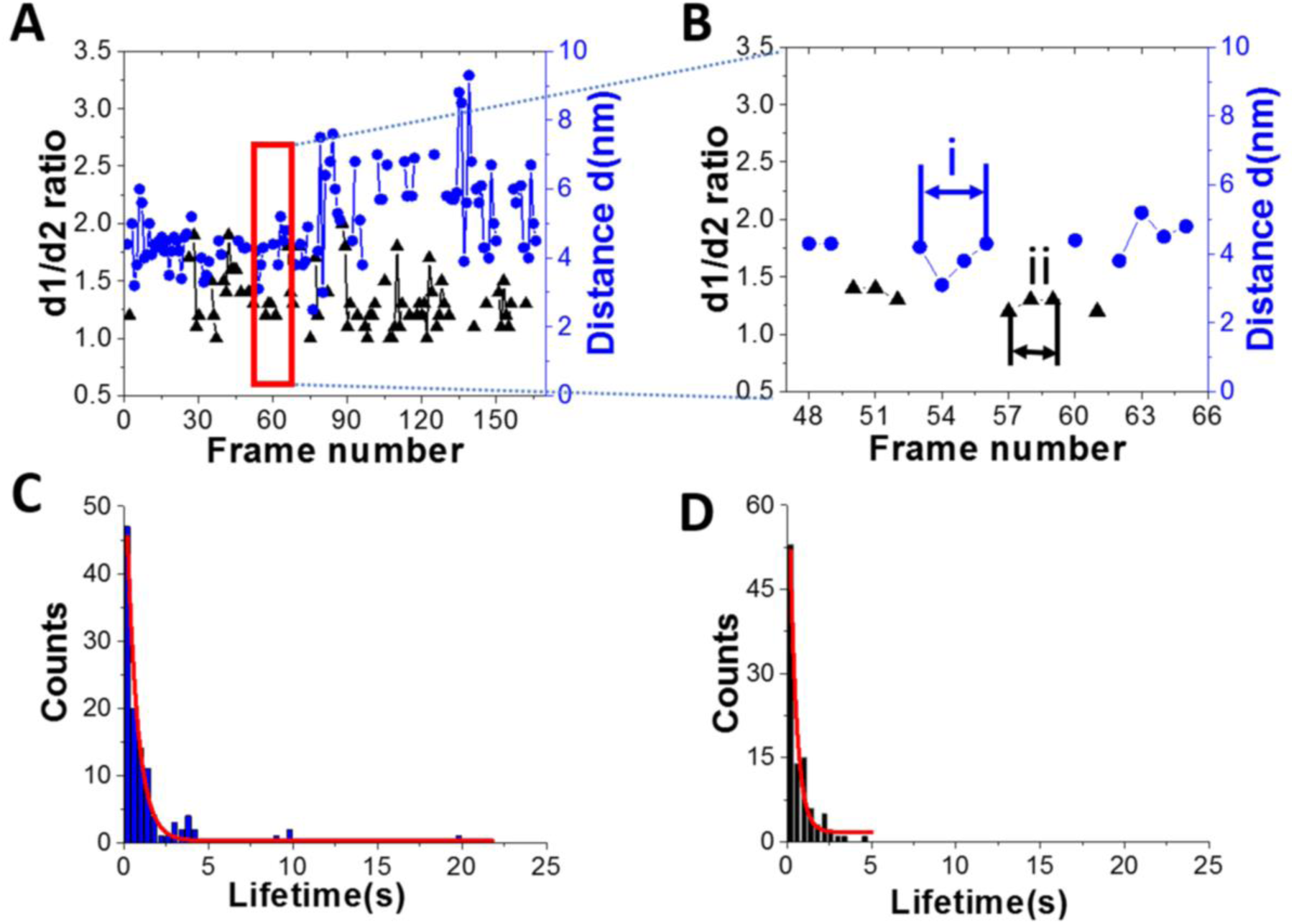
Dynamics of A3G in complexes with 69 nt tail ssDNA. A. Plot illustrating the dynamics of A3G in complex with the 69 nt tail ssDNA. The blue dots represent the distance (d) between two domains for the dumbbell structure of A3G. Black triangles represent the ratio of two orthogonal diameters, d1:d2, for the globular structure of A3G. B. Zoomed view from Figure 4A (marked by red quadrant) for dumbbell (blue dots) and globular (black triangles) structures of A3G. The arrows show examples for calculation of the lifetime for dumbbell structures “i” (blue) and globular “ii” (black) for A3G. C. The histogram for the lifetime of A3G in the dumbbell structure, which after fitting with first-order exponential model, gives a lifetime of ∼0.64 ± 0.03 seconds. D. The histogram for the lifetime of A3G in the globular structure, which after fitting with first-order exponential model gives a lifetime of ∼0.39 ± 0.06 seconds.

### A3G in complex with the 25 nt tail DNA substrate

To understand the role length plays in the ssDNA substrate on the structure and dynamics of A3G, the length of the ssDNA substrate was reduced to 25 nt. The selected frames from Movie 2, as shown in Figure 5, demonstrate the structure and dynamics of monomeric A3G in complex with the 25 nt tail ssDNA. In this complex, A3G also reveals both structures: globular (in frames 1 and 38) and dumbbell (in frames 19 and 35). However, the estimated yield of the dumbbells, calculated from 24 separate movies and ∼600 frames total, was 35%, which is roughly two times less than that of the 69 nt tail ssDNA substrate. Similar to the analysis of the A3G-69 nt tail ssDNA complexes, data was analyzed for the A3G-25 nt tail ssDNA complexes and the results are presented in Figure 6. The dependence of the distance (d) on the frame number illustrates the dynamic properties of the dumbbell structure of A3G, as shown in Figure 6 A. A histogram for the distance (d) is shown in Figure 6 B. For the dumbbell structure of A3G, the mean distance is d = 4.7 ± 1.0 nm, which is slightly less than for the A3G in complex with the 69 nt tail of the ssDNA, which is 5.1 ± 1.0 nm. The results for the globular structure show the dependence of the d1:d2 on the frame number (Figure 6C); the histogram for d1:d2 is shown in Figure 6 D. The calculated lifetimes for dumbbell and globular structures are shown in Figure 6 E and 6 F, which show the lifetime of the dumbbell structure for A3G in complex with the 25 nt tail ssDNA is less than for the globular structure: 0.42 ± 0.01 seconds and 1.29 ± 0.09 seconds, respectively.

**Figure 5.**
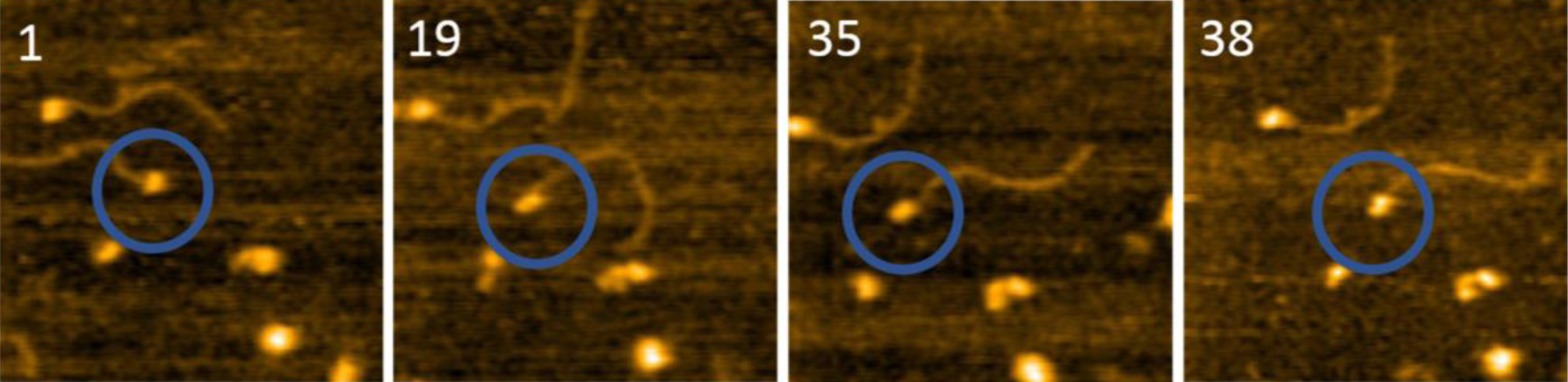
Selected frames illustrating the dynamics of A3G in complex with the 25 nt tail ssDNA. Circles show the complex of the interest. Frames 1 and 35 represent the globular structure of A3G, and frames 19 and 35 show the dumbbell structure. The average yield of dumbbell structures is 35%. The scan size is 200nm, and the scan rate is 398 ms/frame.

**Figure 6.**
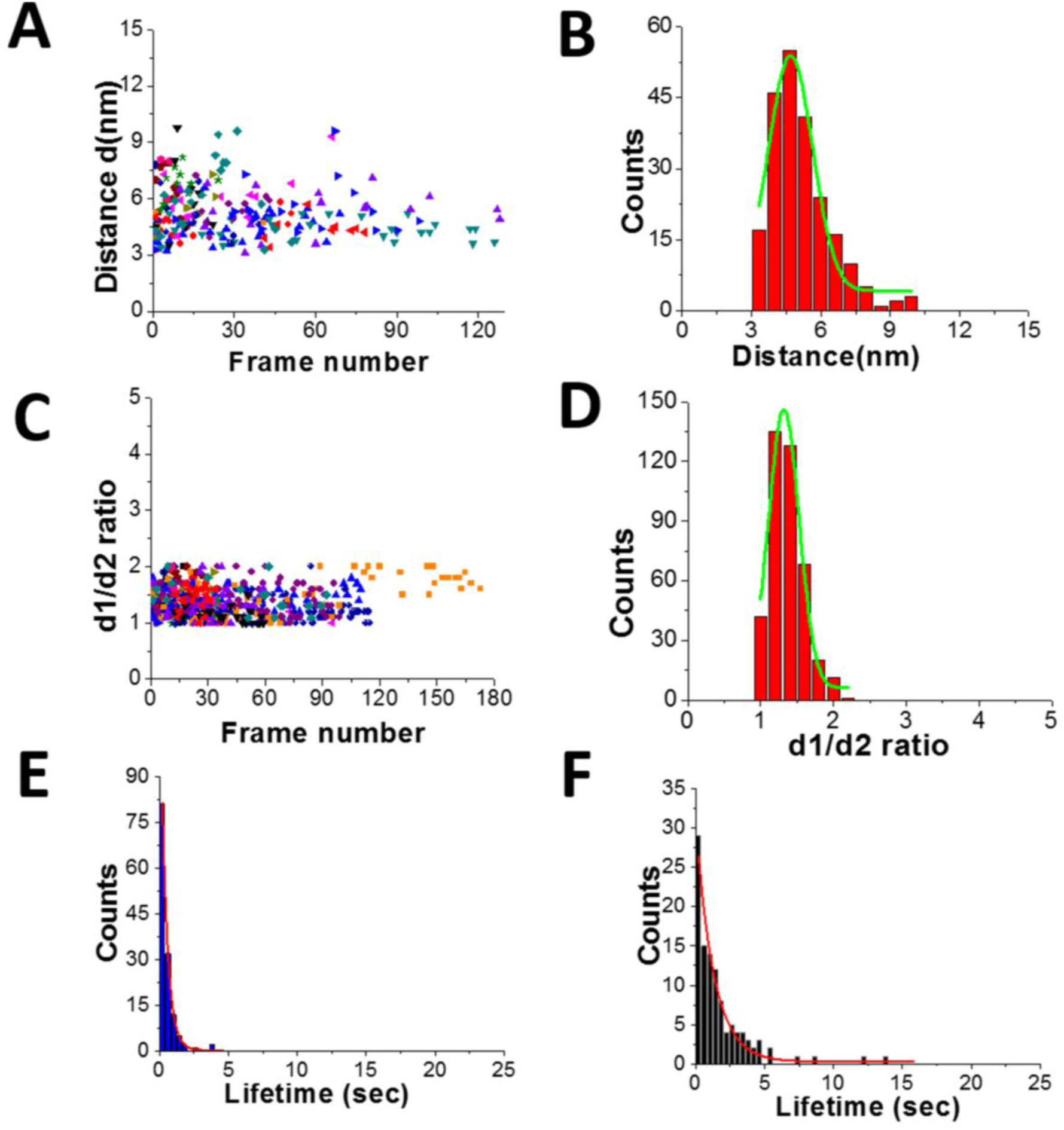
Data analysis for the A3G-25 nt tail ssDNA complex. A. The distance (d) between Domain 1 and Domain 2 for the dumbbell structure of A3G in the A3G-25nt tail ssDNA complex. B. The histogram of the distances between two domains. The mean value for distances between Domain 1 and Domain 2 together with the standard deviation is 4.7 ± 1.0 nm. C. The dependence of d1:d2 ratio for the globular structure of A3G in A3G-25nt tail ssDNA complex on frame number. D. The histogram for the d1:d2 ratio. The mean value for d1:d2, with the standard deviation, is 1.3±0.2. E. The lifetime of dumbbell structure of A3G in complex with the 25 nt tail ssDNA. After fitting, the lifetime is 0.42 ± 0.01 sec. F. The lifetime of the globular structure of A3G in complex with 25 nt tail ssDNA. After fitting, the lifetime is 1.29 ± 0.09 sec. The data are the result of analysis of ∼600 frames from 18 separate movies.

### A3G in complex with the 69 nt gap DNA substrate

The results for the 69 nt tail ssDNA substrate show that the position of one of the domains in the dumbbell structure of A3G changes relative to the dsDNA tag (Figure 1, frames 56, 57). Additionally, one of the domains of A3G appears smaller in size. This observation indicates a possible transient dissociation of one of the domains from the ssDNA substrate. To directly visualize and characterize a possible transient dissociation of one of the domains from ssDNA substrate, the 69 nt gap ssDNA substrate was used, where 69 nt ssDNA was fused between two dsDNA duplexes (Figure S1 C).

Figure 7 presents selected frames from Movie 3, where the transient dissociation of one of the A3G domains from the ssDNA substrate is unambiguously seen. Frames 21, 25, 47, and 56 show one smaller-sized domain unbound to the ssDNA substrate. Frames 175, 182, 187, and 196 show both domains, similar in size, bound to the ssDNA gap substrate. A3G also formed a globular, compact structure, as seen in frames 42 and 73.

**Figure 7.**
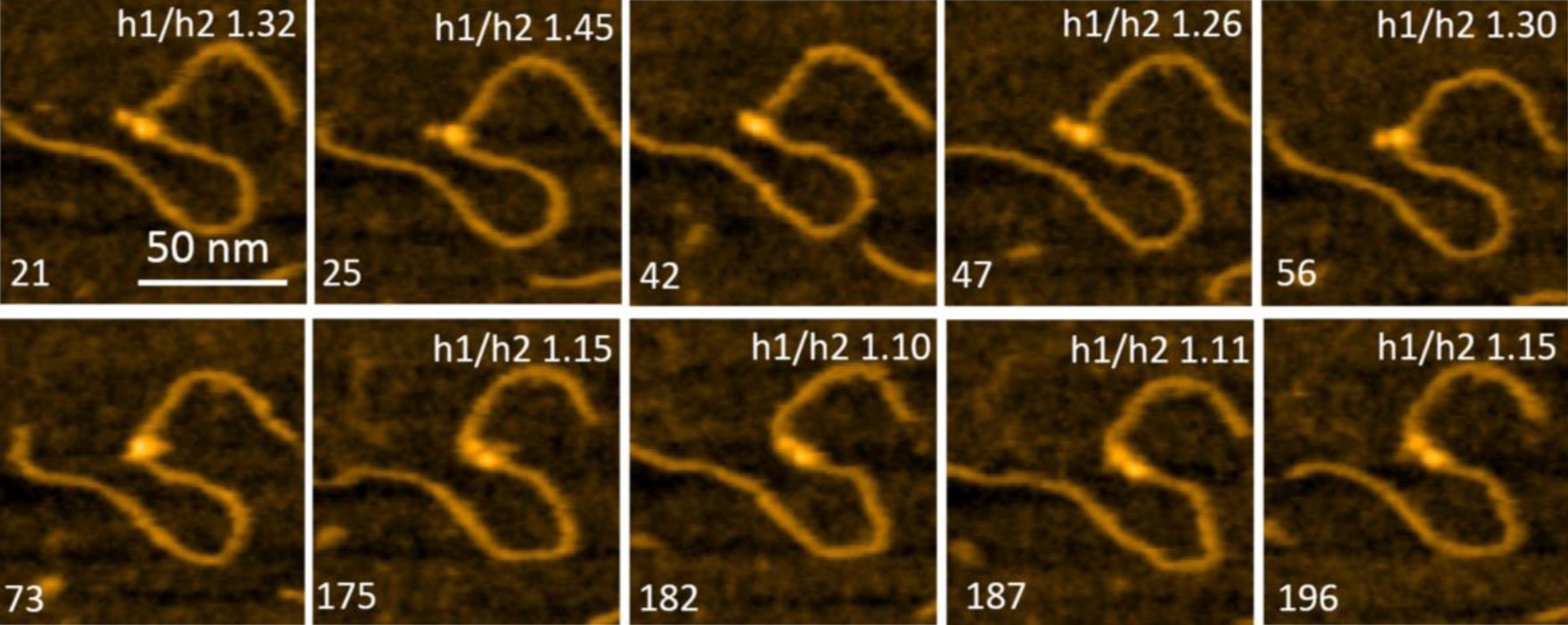
Selected frames from Movie 3 illustrating the different positions of A3G domains in complex with the 69 nt gap DNA. Frames 21, 25, 47, and 56 show the dumbbell structure of A3G with one domain unbound to the ssDNA substrate. Frames 42 and 73 represent the globular structure of A3G. Frames 175, 182, 187, and 196 represent the dumbbell structure of A3G with both domains located on the ssDNA substrate. The ratio of heights of Domain 1 to Domain 2 (h1:h2) of A3G is inserted on the top of each frame. The scale bar is 50 nm. The scan rate is 398 ms/frame.

The smaller size of such a domain can be explained by its lack of binding to the ssDNA substrate, which may contribute to the overall size of the domain. To confirm this effect, the ratios of the heights were calculated for Domain 1 (h1) to Domain 2 (h2) (Figure 2B). Data for the h1:h2 ratio are incorporated into frames in Figure 7. When both domains are in a dumbbell structure and bound to the substrate, DNA contributes equally to the sizes of the domains. Therefore, the ratio of heights of the domains h1:h2 would be expected to be close to one, which is clearly seen in frames 175, 182, 187, and 196. Meanwhile, when one of the domains is unbound to the ssDNA substrate, the h1:h2 ratio should increase due to the lack of binding of this domain with the ssDNA substrate, as seen in frames 21, 25, 26, and 56.

## Discussion

The data presented demonstrate the structure and dynamics of full-length, monomeric A3G in complex with ssDNA substrates. The continuous, frame-by-frame HS-AFM imaging of A3G-ssDNA complexes allowed for not only clear visualization of the dumbbell and globular structures of A3G in complex with ssDNA substrates, but also the transition between them. The major finding here is the high yield of A3G dumbbell structures in complex with ssDNA substrates compared to free protein,^28^ suggesting that the interaction with ssDNA substrates shifts the conformational equilibrium of A3G to the dumbbell conformation.

The yield of the dumbbell conformation of A3G also depends on the length of the ssDNA substrate. Table 1 summarizes data obtained from analyses of the dumbbell and globular structures of A3G in complexes with 69 nt and 25 nt tail ssDNA substrates and free A3G. As seen in Table 1, in the presence of a long, 69 nt ssDNA substrate, the dumbbell structure shows the highest yield of dumbbells (65%), which drops to 35% for a shorter, 25 nt ssDNA substrate, and comprises only 16% for a free A3G. Together these data clearly demonstrate the effect of ssDNA substrate on conformational changes of A3G domains and show the dependence of such changes on the length of ssDNA substrate. The average distance between A3G domains for the dumbbell structures in A3G-ssDNA complexes tends to slightly change, from 5.1 ± 1.0 nm for a long substrate, decreasing up to 4.7 ± 1.0 nm for a shorter one, and being the smallest 4.4 ± 0.9 nm for a free A3G. Data for the globular structure does not demonstrate changes for A3G ssDNA complexes and free A3G, indicating the ssDNA substrate does not affect the globular structure of A3G. Indeed, the d1:d2 ratio remains equal to 1.3, indicating the elongated shape for both, A3G in the complex with ssDNA and free A3G.

**Table 1.**
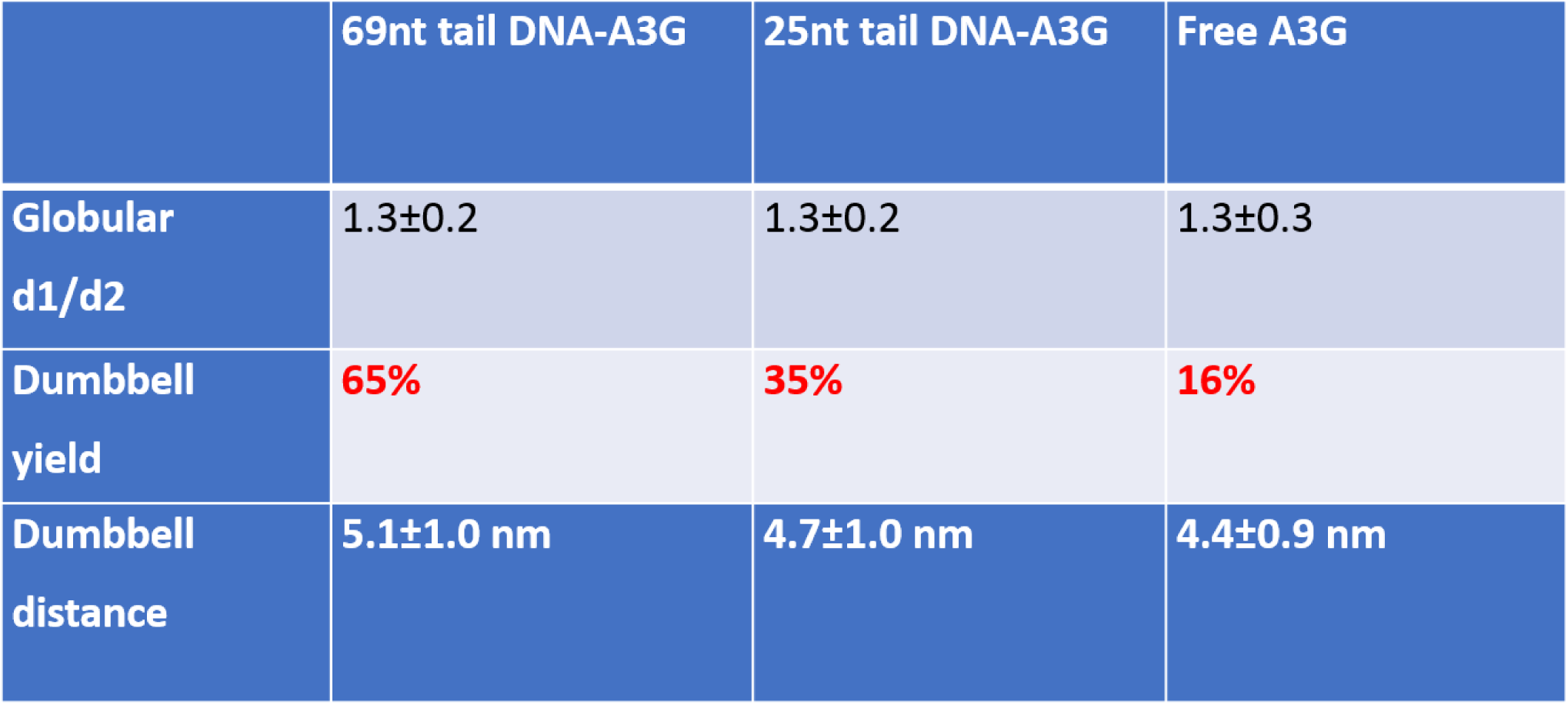
The yield and the distance between domains in the dumbbell structure of free A3G and in complexes with ssDNA.

HS-AFM data also reveal a different affinity for the A3G domains in the dumbbell structure to the DNA substrate. As seen in Figure 7, one of the A3G domains in complex with ssDNA is capable transiently dissociating from the ssDNA substrate. Quantitatively, for the dumbbell structure of A3G in the complex, this effect is illustrated by measuring of ratios of the heights of Domain 1 to Domain 2 (h1: h2). The value of the h1: h2 ratio is close to one when both domains are bound to the substrate, but when one of the domains is unbound to the ssDNA the h1: h2 ratio is 1.3. These measurements were completed for ssDNA substrates with both the 69 nt and 25 nt tails ssDNA. Figure 8A and 8B present the results of this analysis. Histograms for A3G complexes with 69 nt and 25 nt tail ssDNA substrates have two distinct peaks. The first peak, with almost equal heights of the domains, corresponds to cases when both domains are bound to the substrate. The second peak corresponds to cases when one of the domains is unbound to the substrate, with the h1: h2 ratio close to 1.3, indicating on contribution of ssDNA to the size of the domain. Comparatively, for free A3G (Figure S3), the histogram shows only one maximum for the ratio h1: h2, which is close to one. Another line of evidence for the contribution of ssDNA to the overall size of the A3G domains comes from directly measuring the heights of each domain for free A3G and in complex with 69 nt tail ssDNA, as shown in Figure S4. Here, we assembled histograms for the heights of each domain in dumbbell structure for free A3G (Figure S4 A, B) and for A3G in complex with the 69 nt tail ssDNA (Figure S4 C, D). Data demonstrate that the heights of the domains for free A3G are similar when compared to the heights of domains for A3G in the complex (Figure S4 C, D). Note that the height of one of the domains for A3G in the complex with ssDNA substrate is close to the height of both domains for free A3G (Figure S4 D), which indicates this domain unbound to the ssDNA substrate (Figure S4 C). Overall, the data presented here clearly demonstrate that one of the domains in the dumbbell structure of A3G is capable of transiently dissociating from the ssDNA substrate, supported by the lack of the contribution of ssDNA substrate in the size of the protein.

**Figure 8.**
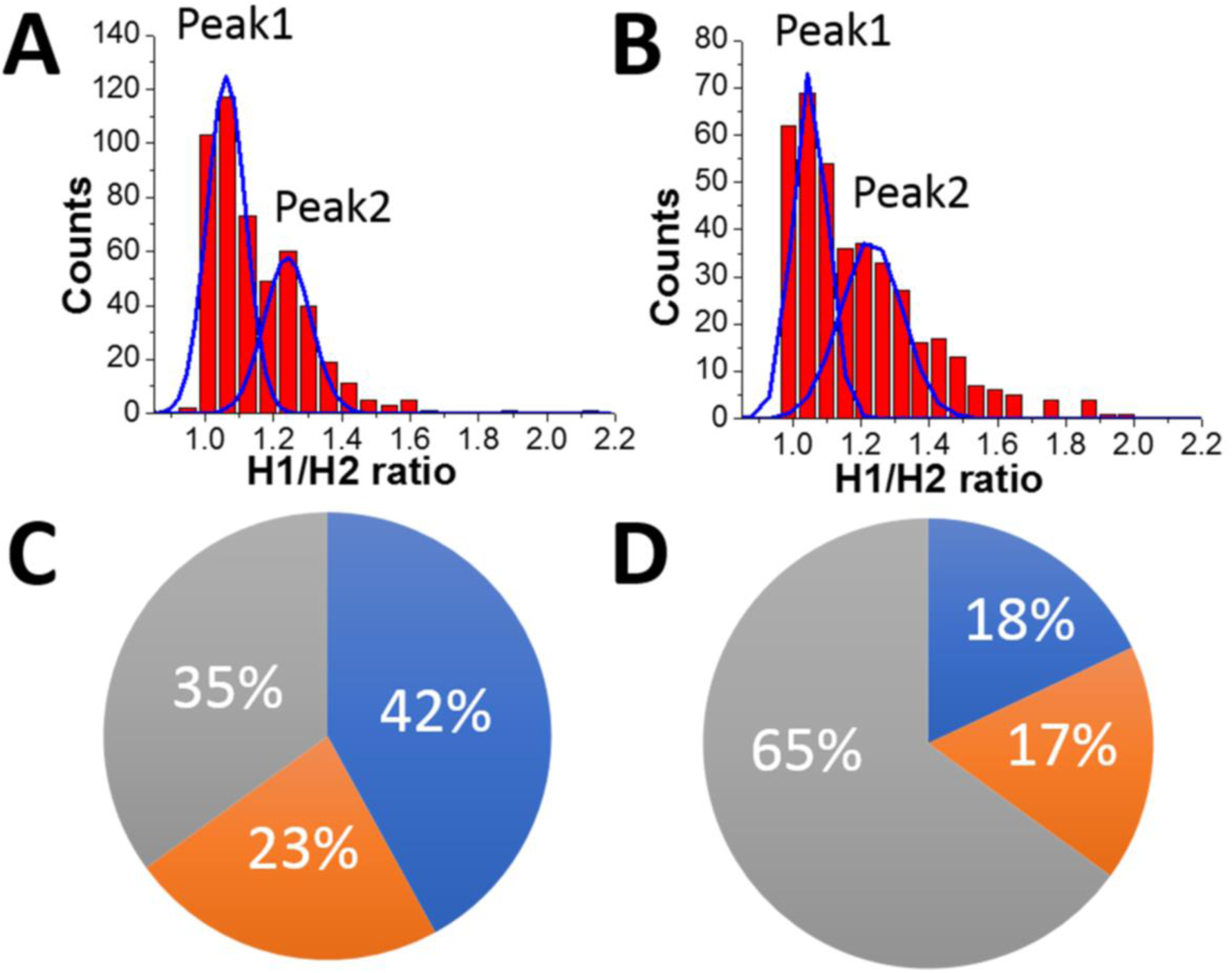
The ratio of the heights of Domain 1 to Domain 2 (h1:h2). A. The A3G-69 nt tail ssDNA complex. B. The A3G-25 nt tail ssDNA complex. The ratios of the areas under the first peak and the second peak is 1.8 for the A3G-69 nt ssDNA complex (A) and 1.1 for the A3G-25 nt ssDNA complex (B). The diagram represents the distribution of globular and dumbbell A3G structures for the 69 nt tail ssDNA substrate (C) and the 25 nt tail ssDNA substrate (D). Grey area shows the yield of globular A3G structures for the 69 nt tail ssDNA substrate with 35% (C) and 65% for the 25 nt tail ssDNA substrate (D). The blue area illustrates both A3G domains bound to the substrate, and the orange area shows one of the domains unbound to the substrate. For long substrates (C), both domains are bound to the substrate in 42% of cases and one domain is unbound from the substrate in 23% of cases. For a short substrate (D), both domains are bound to the substrate in 18% of cases, and one domain is unbound to the substrate in 17% of cases.

The diagram in Figure 8C and 8D summarizes the analysis of all results obtained here. The grey area in the diagram presents the yield of globular A3G structures, calculated as 35% for the 69 nt tail ssDNA substrate (A) and 65% for the 25 nt tail ssDNA substrate (B), respectively. The estimated lifetime for the globular A3G structure in complex with the 69 nt tail ssDNA (∼ 0.39 ± 0.06 sec) tends to be less than for the 25 nt tail ssDNA substrate (∼ 1.29 ± 0.09 sec). The shorter lifetime for the globular structure correlates with the reduced yield of the globular structure compared to the dumbbell structure for the A3G-69 nt ssDNA complexes. The blue and orange areas together show the yield of dumbbell structures for long and short ssDNA substrates as 65% and 35%, respectively, with a tendency toward increased lifetimes for the dumbbell structures in complex with 69 nt ssDNA (∼ 0.64 ± 0.03 sec) compared with a shorter ssDNA substrate (0.42 ± 0.01 sec). These results show the correlation between the yield of dumbbell and globular structures of A3G with their lifetime on different the ssDNA substrates.

As shown in Figure 8A and 8B, the two distinct peaks for h1: h2 values, shown for the long and short ssDNA substrates, demonstrate different positions of A3G domains on the ssDNA substrate. Indeed, when both domains are bound to the ssDNA substrate, the h1:h2 ratio is close to one, compared to the h1:h2 ratio equal to 1.3 when one of the domains unbound to the substrate. In addition to the position of the domains in dumbbell structures of A3G in A3G-ssDNA complexes discussed above, the areas under peak 1 and peak 2 (Figure 8A and 8B) indicate the different number of events for bound and unbound domains for long and short ssDNA substrates. Indeed, for a long substrate, the ratio of areas under peak 1 and peak 2 is 1.8, indicating on almost twice greater number of events when both A3G domains are positioned on the ssDNA compared to one of the domains being unbound. The blue and orange areas in Figure 8C illustrate such distribution as 42% for both domains bound to the ssDNA substrate (blue area) vs 23% for the unbound one (orange area). For a short ssDNA substrate (Figure 8D), the ratio of areas under peak 1 to peak 2 is 1.1, demonstrating a practically equal number of events for A3G domains positioned on the substrate and one domain unbound, as shown in blue (18%) and orange (17%) areas in the diagram, respectively.

HS-AFM is not capable of identifying which domain remains in contact with the ssDNA and which is temporary dissociated. Nevertheless, several lines of evidence allow us to posit the CTD is the domain capable of transiently dissociating from the ssDNA. Computer analysis performed^35^ shows the isoelectric point (pI) of the N-terminal domain (NTD) is 9.6, compared to 6.9 for the CTD. In addition, the number of aromatic amino acids in A3G essential for ssDNA binding is 9 for the NTD versus only 6 for the CTD. Taken together, these findings suggest tighter binding for the NTD than for the CTD. Also, more stable binding of NTD with ssDNA than CTD has been reported.^28,35,36^ Moreover, it is demonstrated that the NTD is responsible not only for binding with ssDNA,^35,36^ but for positioning and stabilizing active sites of the CTD for efficient deamination of ssDNA.^37^ Mutational studies^38^ suggest the following two steps for A3G binding with ssDNA template: (1) initially, high affinity contacts are carried out by the NTD with Kd in the nM range, (2) followed by the CTD with Kd in the µM range. In addition, the data obtained in.^39,40^ have demonstrated that during A3G sliding, the CTD tends to dissociate from ssDNA Therefore, we hypothesize that the CTD has greater conformational mobility compared to the NTD, and is capable of transiently dissociating from ssDNA template.

Based on our data, we suggest a model where the substrate length is key in determining whether a dumbbell or globular structure will form on each ssDNA substrate. Figure 9 illustrates such a model for a long (A) and a short (B) ssDNA substrates. The red ball represents CTD, which forms a dumbbell structure and is unbound to the ssDNA, and the blue ball represents the NTD bound to ssDNA (state i). In this state (i), only the NTD is bound to the substrate, and A3G may dissociate from a long or short substrate with an equal probability. This would explain the similar number of cases when only one domain is bound to the substrate for both long and short ssDNA substrates, 23% vs 17%, respectively (Figure 8C, D orange area). If not dissociated, as in the case of a long substrate (A), the CTD may return to the substrate and preserve the dumbbell structure (grey arrows, states ii) with both domains bound to ssDNA; alternatively, A3G may come close to the NTD domain to form a globular structure (purple arrow, state iii). In the case of a long substrate, A3G has a greater chance of holding the dumbbell structure with both domains bound to the ssDNA, as shown in Figure 8C (blue area). Therefore, it is reasonable to theorize that for a long substrate, the increased yield of dumbbell structures is primarily due to both domains being bound to the substrate. However, this differs for a short substrate (B). Indeed, the CTD in state (i) may return to the NTD domain to form a globular shape (purple arrow, state iii) or form a preserved dumbbell structure with both domains bound to the substrate (grey arrow, state ii) or one domain dissociated from the substrate (orange arrow, state iv). However, for a short substrate, there is less possibility to preserve the dumbbell structure with two domains bound to the substrate, which comprise 18% (Figure 8D blue area), compared to 42% for a long substrate (Figure 8C, blue area).

**Figure 9.**
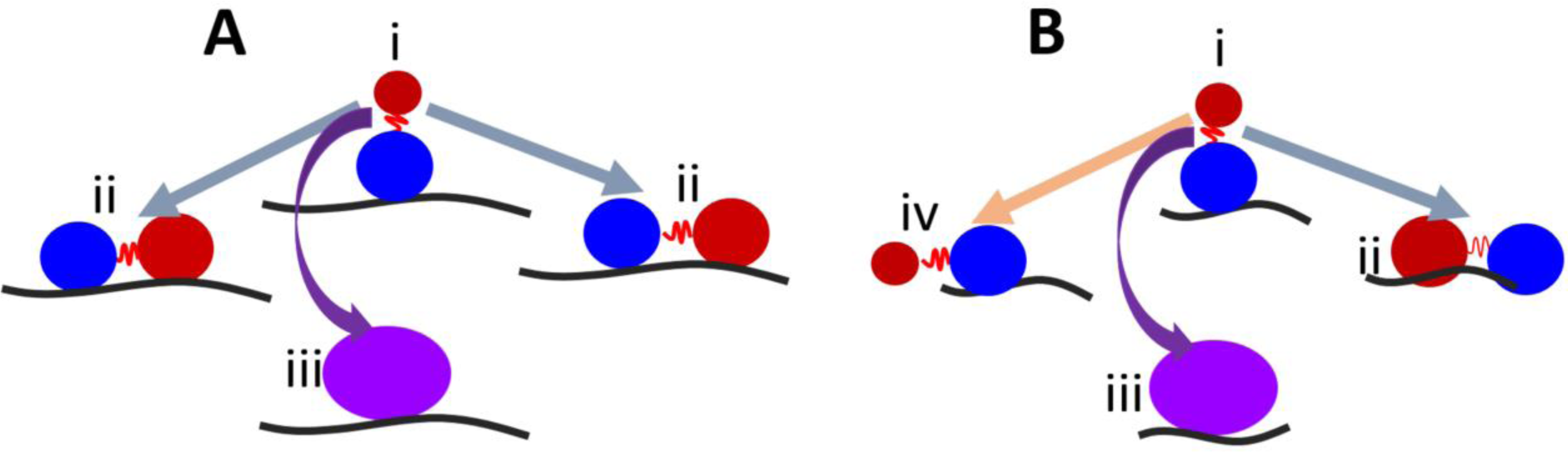
The model explaining the role of the dumbbell conformation of A3G in the assembly complexes with ssDNA. The red and blue balls represent the CTD and NTD, respectively. State i illustrates the dumbbell structure with one A3G domain unbound to the ssDNA substrate. In state i, A3G is capable of transiently dissociating/associating from the substrate. In state ii, both domains are bound to the ssDNA substrate (grey arrows) or form a globular structure (iii) bound to the ssDNA substrate (purple arrow). State (iv) shows one domain unbound in case of short substrate (orange arrow).

The conformational changes between domains, provided by an interdomain linker,^28^ are easier achieved when A3G adopts a dumbbell structure and may facilitate functions of A3G such as sliding,^8, 19,41^ intersegmental transfer,^16^ and eventually the search for deamination of the ssDNA substrate. Our data demonstrate that one of the domains is capable of transiently dissociating from the substrate, and such a dynamic may facilitate the search for the deamination target. Moreover, we suggest CTD is the domain that transiently dissociates from the substrate to facilitate this search. Based on our data, we posit the dumbbell structure of A3G represents an active structure of the protein. Interestingly, a drop in yield for dumbbell structures with a short substrate correlates with the length dependence of deaminase activity of A3G.^8,42,43^ Indeed, it was shown^8^ that specific activity of A3G increases between 15 nt and 60 nt ssDNA lengths and remains unchanged thereafter. Despite that both globular and dumbbell forms of A3G provide efficient binding with the ssDNA substrate, a correlation between length-dependence of deaminase activity and the yield of dumbbells support our hypothesis that dumbbell structures of A3G represent an active form of the protein.

## Materials and Methods

### Hybrid ssDNA substrates

#### The 69 nt tail ssDNA

The hybrid 69 nt tail ssDNA was assembled as previously described.^33^ Briefly, the synthesized (Integrated DNA Technology; IA) 89 nt oligo was annealed at a 1:1 ratio with phosphorylated 23 nt oligo (Integrated DNA Technology, IA) to form a 20 bp DNA duplex with sticky ends. Later, the construct was ligated at 16°C overnight with a previously gel-purified 356 bp DNA fragment with sticky ends. The ligated product was purified from the gel using QIA quick Gel Extraction Kit (Qiagen) as described,^33^ and re-suspended in TE buffer containing 10mM Tris, pH 7.5, and 1mM EDTA. The final product consists of the 69 nt ssDNA attached to a 379 bp dsDNA fragment as a tag.

#### The 25nt tail ssDNA

The hybrid 25 nt tail ssDNA was assembled according to the schematics described directly above for 69 nt tail ssDNA. In this case, synthesized 58 nt oligos (Integrated DNA Technology, IA) were annealed with phosphorylated 20 nt oligos to create a 33 bp duplex with a sticky end to ligate with a 224 bp DNA fragment. The final product consists of the 25 nt tail ssDNA attached to a 260 bp dsDNA as a tag.

#### The 69 nt gap ssDNA

Creation of the hybrid DNA substrate, in which an ssDNA region is flanked by dsDNA arms, has been previously described in detail.^44,45^ First, 235 bp dsDNA and 441 bp dsDNA fragments with sticky ends were generated by PCR and purified from the gel. Second, 235 bp hybrid 5’end tail ssDNA and 441 bp hybrid 3’ end tail ssDNA substrates were prepared as described above for preparation of hybrid tail ssDNA substrates. Third, two hybrid 3’ and 5’ end tail ssDNA substrates were mixed at a 1:1 ratio and annealed with the bridge oligo. Next, the annealed product was ligated at 16°C overnight. To remove the bridge oligo, the product was heated to 70^°^C for 5 minutes and immediately put into ice. Finally, the 69 nt gap DNA substrate was gel purified by QIAquick Gel Extraction Kit (Qiagen), as described.^33^ The final product consists of 69 nt ssDNA flanked with 441 bp and 235 bp dsDNA, respectively.

### Preparation of A3G in complex with ssDNA substrates

For each ssDNA substrate mentioned above, the complex with A3G was formed at a 4:1 protein-to-ssDNA ratio in binding buffer containing 50 mM HEPES, pH 7.5, 100 mM NaCl, 5 mM MgCl2, and 1 mM DTT. The complex was incubated for 15 minutes at 37°C before deposition on a mica surface. Figure S1 schematically shows the positions of A3G on different ssDNA substrates

### Sample preparation for HS-AFM

A detailed description of the sample preparation for HS-AFM has been previously described.^17^ In brief, a small piece of mica, glued to the cylinder, was cleaved and treated with APS, as described.^17^ Two microliters of the complexes were deposited on the APS mica surface for 2 minutes, followed by washing with binding buffer. Continuous scanning was initiated immediately following the wash, without drying of the sample. The selected scanning area (200 nm x 200 nm) was continuously imaged to visualize the dynamics of the complexes at a scan rate of 398 ms/frame. The tips for imaging were grown under an electron beam using short cantilevers (BL-AC10DS-A2, Olympus; Tokyo, Japan) with spring constant between 0.1 – 0.2 N/m and a resonance frequency of 400 – 1000 kHz.

### Analysis of the HS-AFM data

After collecting frame-by-frame HS-AFM images for A3G in complex with different ssDNA substrates, a set of movies was assembled. Analysis of these movies revealed the following two structures for A3G in the complexes: dumbbell and globular. To analyze the data obtained from HS-AFM experiments, the cross-sectional feature was used in FemtoScan Online software (Advance Technologies Center; Moscow, Russia), as previously described.^28,33,46^ Analysis was completed for each frame from the collected movies. More than 500 frames were analyzed for each A3G structure in the A3G-ssDNA complexes.

## Author Contributions

Y.L., L.S, and Y.P equally planned the research and contributed to the writing of the manuscript. Y.P performed the experiments, and L.S and Y.P analyzed the data and designed the figures.

## Acknowledgments

This work was supported by NIH grants awarded to Dr. Lyubchenko (R01-GM118006, GM096039, and R21NS101504). We thank Dr. R. Harris for providing A3G protein and Dr. Lyubchenko’s lab members for their fruitful discussions. We thank Mohtadin Hashemi for his productive and helpful discussions regarding this work and Dr. Atsushi Miyagi for HS AFM movie of 69nt gap ssDNA-A3G complex. Lastly, we thank Melody A. Montgomery for the professional editing of this manuscript.

## Table of contents

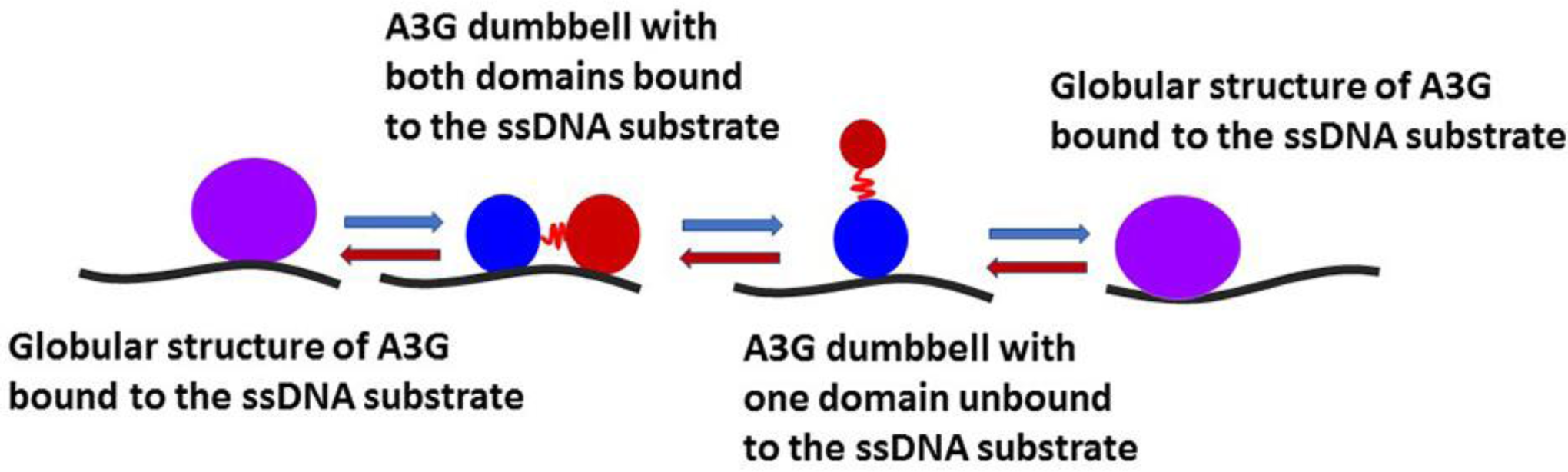

