## Supporting information for "Insight into dynamics of APOBEC3G protein in complexes with DNA assessed by high speed AFM"

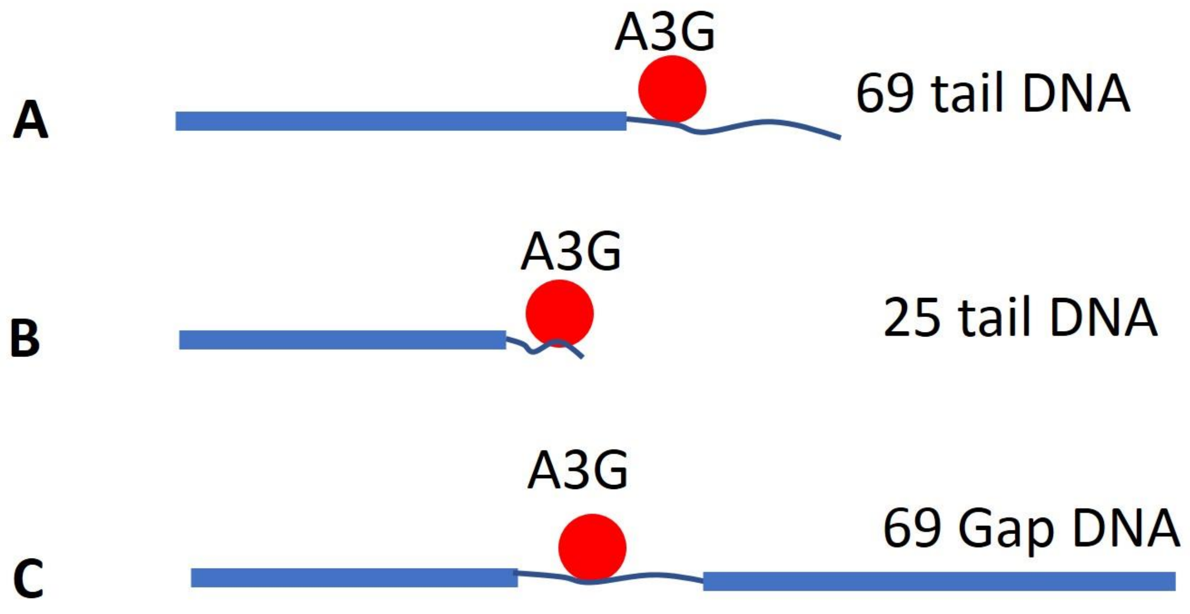


**Figure S1. Schematic presentation of tails and gap hybrid DNA constructs. A.** The 69 nt tail ssDNA hybrid. **B.** The 25 nt tail ssDNA hybrid. **C.** The 69 nt gap ssDNA hybrid. Thick blue lines correspond to the dsDNA tags used for clear identification of the complexes by AFM. Red balls represent A3G protein attached to the ssDNA part (thin blue line). For tails DNA substrates, A3G appeared as a blob at the end of dsDNA tags. For the A3G-gap DNA complex, A3G is seen in the middle of two dsDNA tags.

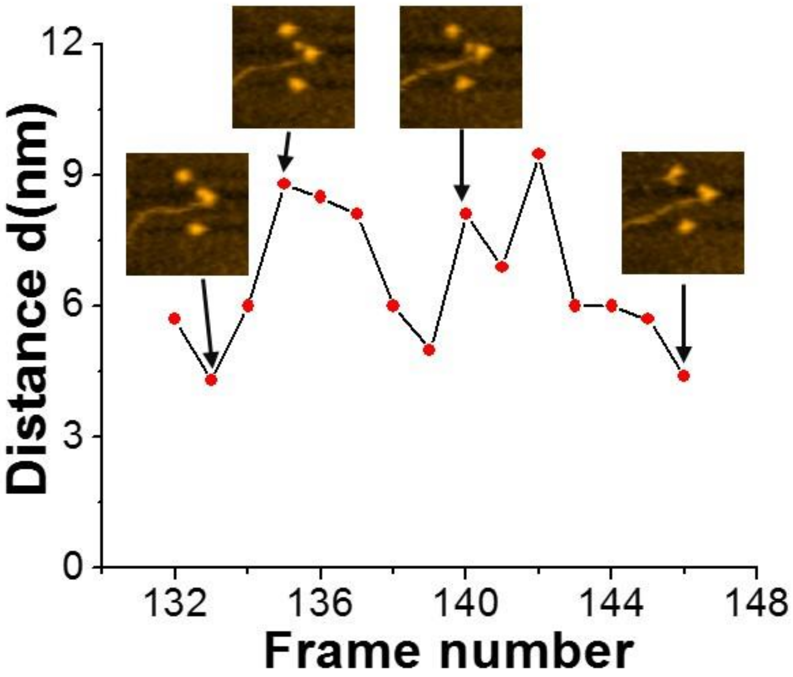


**Figure S2. Plot showing the dynamics of the dumbbell structure of A3G in complex with 69 nt ssDNA complex.** The arrows point to the maximum and minimum distances between A3G domains, measured from the corresponding AFM frames.

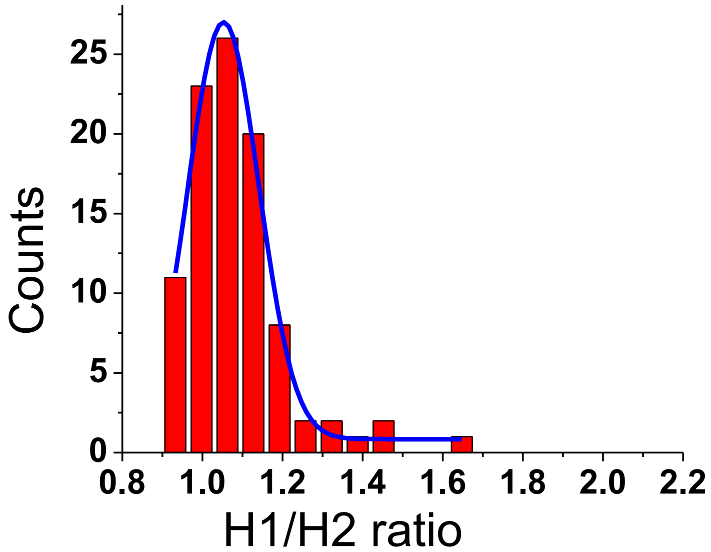


**Figure S3. The ratio of heights for Domain 1 (h1) and 2 (h2) for free monomeric A3G.**

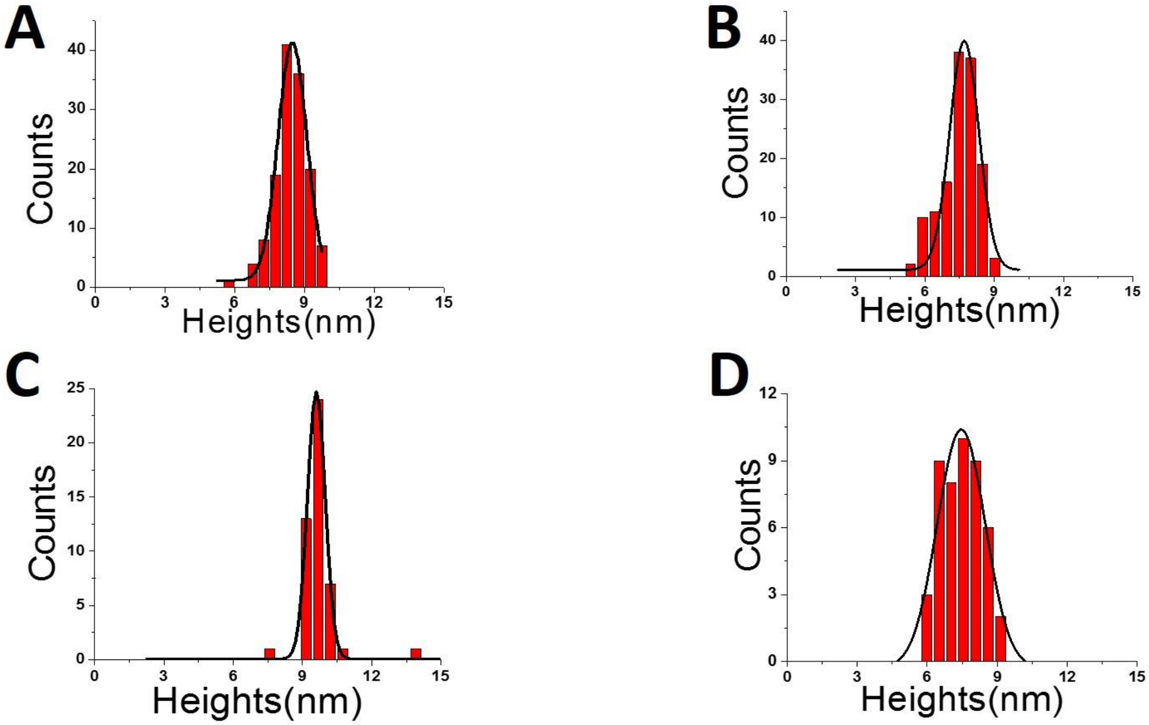


**Figure S4. The histograms of heights for Domains 1 and 2 for dumbbell structures of free A3G (A, B) and in complex with 69nt ssDNA substrate (C, D).** The values for the heights are shown. All measurements are performed from the same AFM image to exclude possible measurement error.
